# Inhibitory control, personality, and manipulated ecological conditions influence foraging plasticity in the great tit

**DOI:** 10.1101/2020.12.16.423008

**Authors:** Jenny R. Coomes, Gabrielle L. Davidson, Michael S. Reichert, Ipek G. Kulahci, Camille A. Troisi, John L. Quinn

## Abstract

1. Organisms are consistently under selection to respond effectively to a diversity of, sometimes rapid, changes in their environment. Behavioural plasticity can allow individuals to do so instantaneously, but why individuals vary in this respect is poorly understood. Although personality and cognitive traits are often hypothesised to influence plasticity, the effects reported are highly inconsistent, which we hypothesise is because ecological context is usually not considered.
2. Here we explore the roles of individual cognitive and personality variation – assayed using standard tasks for inhibitory control, a measure of self-control, and ‘reactive-proactive’ personality axis (RPPA), respectively – in driving foraging plasticity, and asked how these effects varied across two experimentally manipulated ecological contexts: food value and predation risk.
3. After great tits (*Parus major*) had initially been trained to retrieve high value food hidden in sand, they were then simultaneously offered the hidden food and an alternative food choice on the surface, that was either high or low value. Their choices were further examined under high and low perceived predation risk treatments. Individuals’ choices were classified in terms of whether they continued to forage on the hidden but familiar food source, or instead switched to the new visible food source. We defined the latter option as the plastic response.
4. Our assays captured consistent differences among individuals in foraging behaviour. Both inhibitory control and exploration influenced whether birds switched from the familiar but hidden food source to the new alternative visible food on the surface. These effects depended on the relative value of the food items available and on the perceived level of predation risk, but also on the time scale over which the response was measured.
5. Our results demonstrate how an executive cognitive function and one specific personality axis can simultaneously influence plasticity in a key functional behaviour. That their effects on foraging were primarily observed as interactions with food value or predation risk treatments also suggest that the population level consequences of behavioural mechanisms, such as these, may only be revealed across key ecological conditions or gradients.

## Introduction

Organisms are consistently under pressure to adapt to changes in their environment, such as, changes in climate, food availability, and predation risk. Behavioural plasticity allows individuals to respond to environmental change by adjusting their behaviour (Gross, Pasinelli and Kunc, 2010; Snell-Rood, 2013), but behavioural plasticity is constrained by the costs of sampling information (Dall and Johnstone, 2002; Snell-Rood, 2013) and adjusting behaviour (Komers, 1997). Although these costs are ubiquitous, some individuals are more plastic than others (Dingemanse *et al*., 2009; Coppens, De Boer and Koolhaas, 2010). Why individuals vary in their plasticity is a major focus of research in evolutionary ecological studies of behaviour (Wolf, Van Doorn and Weissing, 2008; Dingemanse *et al*., 2009). Cognition and personality have the potential to influence behavioural plasticity (Dingemanse *et al*., 2009; Snell-Rood, 2013) but their role in doing so under realistic ecological scenarios is poorly understood.

A major focus in cognitive ecology has been to explore the cognitive traits that influence fitness (Cole *et al*., 2012; Shaw *et al*., 2019; Sonnenberg *et al*., 2019), and in particular, foraging success (Balda and Kamil, 1992; Healy and Hurly, 1995). Most of the focus has been on the role of learning, memory, or innovation (Laland and Reader, 1999; Raine and Chittka, 2008; Morand-Ferron *et al*., 2011). Inhibitory control, an executive cognitive function defined as the suppression of a dominant (prepotent) behaviour in favour of a more beneficial or appropriate behaviour (Diamond, 2013), is likely also an important mechanism because foraging success relies on making optimal choices among different options. One study demonstrated that cotton-top tamarins discount future high-quality rewards in favour of immediate lower-quality rewards, a strategy that may be beneficial given the temporal availability of their natural diet (Stevens *et al*., 2005). Other studies show that inhibitory control predicts dietary breath, although the direction of this relationship varies; primate species with higher self-control had greater dietary breath (MacLean *et al*., 2014), while pheasants with greater dietary breadth had poor performance on inhibitory control tasks (van Horik *et al*., 2018). In general, however, there is a dearth of empirical evidence to support the expectation that inhibitory control influences foraging decisions in ecologically-relevant contexts.

In addition, cognitive performance can vary dramatically depending on the specific conditions (Cauchoix, Chaine and Barragan-Jason, 2020). In particular, the risk of predation may influence the extent to which individuals can suppress the prepotent response and choose an alternative, more rewarding behaviour (Schwabe and Wolf, 2009), as stress in humans promotes habitual behaviours and reduces goal-directed behaviour (Schwabe and Wolf, 2009). Similarly, in the presence of a predator, individuals might be expected to minimize predation risk rather than expending time and resources inhibiting a prepotent response. Therefore, an individuals’ “ability” to perform alternative behaviours may not only be dependent on their inhibitory control abilities, but also on the environmental context: when under predation risk, individuals may focus on avoiding predation and show little variation in behavioural plasticity, and when not under predation risk, individuals may focus on the task at hand, and show variation in behavioural plasticity (MacLean *et al*., 2014; Rosati, 2017). Nevertheless, the effects of inhibitory control on behavioural plasticity remain poorly understood, especially when individuals are under predation risk.

Personality refers to consistent differences between individuals in behaviour or behavioural correlations (Sih *et al*., 2004), and is an increasingly common paradigm for examining the evolutionary ecology of behaviour and constraints on behavioural plasticity (Dingemanse *et al*., 2009; Herborn *et al*., 2014). The ‘reactive-proactive personality axis’ (RPPA), for example, contrasts ‘proactive’ individuals, who are more exploratory and risk-prone at one end of the continuum, with ‘reactive’ individuals at the other end, who are less exploratory and more risk-averse (Groothuis and Carere, 2005; Réale *et al*., 2007). Two contrasting hypotheses can explain how the reactive-proactive axis might influence individual behavioural plasticity when conditions change (Arvidsson and Matthysen, 2016; Rojas-Ferrer, Thompson and Morand-Ferron, 2019). The information-gathering strategy (IGS) hypothesis posits that individuals vary in how they collect information from the environment: proactive individuals explore their environment more and sample in novel areas, while reactive individuals explore less and sample known areas (Arvidsson and Matthysen, 2016; Rojas-Ferrer, Thompson and Morand-Ferron, 2019). This leads to the expectation that proactive individuals should display greater behavioural plasticity than reactive individuals, when ecological conditions change. In contrast, the behavioural flexibility (BF) hypothesis states that proactive individuals are more routine-like in their behaviour (Arvidsson and Matthysen, 2016) and are less responsive to changes in their environment (Coppens, De Boer and Koolhaas, 2010); so for example, proactive individuals are less plastic in their behaviour than reactive individuals when faced with a depleted food patch (Verbeek, Drent and Wiepkema, 1994). The conditions under which these divergent predictions are supported are poorly known.

Our aim was to investigate whether inhibitory control and the reactive-proactive personality axis influenced foraging plasticity in a realistic scenario, and whether these effects varied depending on the relative value of alternative food options, and perceived predation risk. We tested this in great tits (*Parus major*), a model species for studies on individual variation in cognition (Cole, Cram and Quinn, 2011; Amy, van Oers and Naguib, 2012; Morand-Ferron *et al*., 2015) and personality (Verbeek, Drent and Wiepkema, 1994; Marchetti and Drent, 2000; Dingemanse *et al*., 2012). First, we performed standard assays for inhibitory control and the RPPA. Next, we trained individuals to retrieve hidden, patchy, high-value food underneath sand, and examined how cognition and personality affected whether individuals continued to use this foraging strategy or instead switched to an alternative, more obvious food source introduced on the surface, while manipulating two variables: 1) the value of the alternative food source, and 2) the risk of predation. Rather than predetermine the adaptive value of each choice, which is difficult to quantify because of context and state dependency, we simply considered the sand or surface food options as alternative choices that were freely available to all individuals.

Our prediction for how inhibitory control could influence whether individuals switched to the visible food source varied depending on which of the choices became the prepotent (dominant) response. In realistic ecological scenarios, the prepotent response is difficult to predict due to conflicts between how the brain simultaneously processes information from past and present stimuli (Anderson and Weaver, 2009). On the one hand, the prepotent behaviour could be to continue the sand foraging technique the birds had been trained to do. In this case, we expected individuals with poor inhibitory control, as measured by the detour-reaching task, to continue to search for hidden food items, even when there were similar food items on the surface (see Table 1). We also predicted in this case that individuals with good inhibitory control could suppress their prepotent foraging technique and instead choose the visible food item when it was of similar value to the hidden food. On the other hand, if the prepotent behaviour is to immediately forage on visible food items, then we expected individuals with poor inhibitory control to feed on the visible food, even when of lower value than the hidden food. Additionally, individuals with good inhibitory control should be able to resist the prepotent response to the visible food when it is low value and instead continue to search for the hidden food. When the visible food is of similar value to the hidden food however, all individuals are likely to choose the visible food.

**Table 1.** Predictions of how inhibitory control (IC), as measured from the detour-reaching task, and personality, as measured from the ‘reactive-proactive personality axis’ (RPPA), influence whether individuals switch from feeding on the hidden high-value food to feeding on the visible low-value (scenario 1) or high-value (scenario 2) surface food. Note that for illustrative purposes, our continuous measure of IC has been changed to a binary ‘good’ or ‘poor’. We refer to visible food choices as representing a plastic response (i.e. a switch) relative to their trained behaviour of foraging in the sand.

|  |  | Food choice |  |
| --- | --- | --- | --- |
|  |  | Scenario 1 | Scenario 2 |
|  | Individual phenotype | Visible low-value, hidden high-value | Visible high-value, hidden high-value |
| a) IC hypotheses |  |  |  |
| i) Hidden food is the prepotent response | Good IC | Choose hidden | Switch to visible |
|  | Poor IC | Choose hidden | Choose hidden |
| ii) Visible food is the prepotent response | Good IC | Choose hidden | Switch to visible |
|  | Poor IC | Switch to visible | Switch to visible |
| b) RPPA hypothesis |  |  |  |
| i) Behavioural flexibility | Proactive | Choose hidden | Choose hidden |
|  | Reactive | Choose hidden | Switch to visible |
| ii) Information gathering | Proactive | Switch to visible | Switch to visible |
|  | Reactive | Choose hidden | Choose hidden |

We predicted personality could also influence foraging plasticity in one of two contrasting ways (Table 1). In line with the IGS hypothesis (Arvidsson and Matthysen, 2016; Rojas-Ferrer, Thompson and Morand-Ferron, 2019), we expected proactive birds would be more likely to switch to an alternative, novel, visible food source, primarily when the alternative food source was of high value, whereas reactive birds would be less likely to utilize the alternative food. Alternatively, according to the BF hypothesis, we predicted reactive birds to be more responsive to the sudden availability of a new food source and to switch to the alternative visible food, especially when it was high value. With the high value food on the surface, there would no longer be a trade-off between food value and searching time. Additionally, we expected the proactive birds to continue foraging on the hidden food source irrespective of the value of alternatives. Finally, given the expectation that the influence of inhibitory control and the RPPA could be context dependent (Stevens *et al*., 2005; Sih and Del Giudice, 2012; Tsukayama, Duckworth and Kim, 2012; Bray, Maclean and Hare, 2014), and that individual differences in behaviour are sometimes only exposed under stress (Suomi, 2004; Quinn and Cresswell, 2005), we explored whether the association between foraging plasticity and inhibitory control, and between foraging plasticity and exploration behaviour, varied depending on predation risk. We expected the great tits, under predation risk, to perform their trained behaviour of searching for the hidden food. To demonstrate the evolutionary validity of our measure of foraging, we also estimated the repeatability of food choices across treatments, which sets the upper limit of heritability, and examined whether any observed between individual variation changed after controlling for potentially confounding effects of the main treatments, or by inhibitory control and exploration.

## Materials and methods

### Aviary housing

We caught wild great tits at seven field sites (three mixed deciduous and four coniferous) in County Cork, Ireland and held them in the aviary on the university campus for a maximum of two weeks from January to March 2018. We fitted birds with a colour ring and a British Trust for Ornithology ring for identification, before placing them in individual cages (62 × 50 × 60cm, H × W × D). When not participating in experiments, birds were fed ad-libitum sunflower seeds, peanuts and water with added vitamin drops (AviMix^®^). Mealworms (*Tenebrio molitor*) were provided three times a day and during experimental training and tests. Before each experiment, we deprived birds of food, but not water, for one hour.

### Exploration assay and inhibitory control assays

On the day after their arrival to the aviary, we released the birds into an experimental room (4.60 × 3.10 × 2.65m, W × L × H) to run the open field ‘exploration of a novel environment’ assay (Verbeek, Drent and Wiepkema, 1994). The experimental room was adjacent to the birds’ individual cages and had five artificial trees (1.53m tall) spaced two metres apart from one another. The number of hops and flights made on the ‘trees’ within two minutes of entering the room was totalled to give each bird an ‘exploration’ score.

On the following day, we assayed inhibitory control using a detour-reaching task in the individual cages, following the methods described in MacLean *et al*. (2014). The detour-reaching task involved presenting a plastic cylinder (3.5 × 3cm, D × L) laterally to the bird, 20 cm in front of a perch that was 5 cm high, so that the bird was positioned in the middle of the long edge of the cylinder before making an approach towards the cylinder. The assay had three phases: 1) Habituation – the birds had to acquire half a waxworm (*Galleria mellonella*) from the open end of an opaque cylinder three times; 2) Training – half a waxworm was placed in the centre of the cylinder and to complete training, birds had to retrieve the food without pecking at the cylinder, in four out of five consecutive attempts; and 3) Test – the opaque cylinder was replaced with a transparent cylinder, and birds were given 10 trials to attempt to retrieve half a waxworm from the centre. During the test phase, any contact a bird had with the cylinder was scored as a fail, and following a failure, the cylinder was removed from the cage. Birds that pecked at the barrier could still access the reward (>90% of failed trials resulted in the bird immediately moving to the side to access the worm). A successful trial was when the bird moved around to the side of the tube and took the waxworm from the open end, as in training. The birds’ final score was the proportion of trials that were successful i.e., high values indicate high inhibitory control (Davidson G.L. unpublished data).

### Experiment pre-training and training

We gave the birds a food preference test consisting of three mealworms and three dehusked sunflower seeds, and recorded the first food they ate. Four individuals did not choose either food in 5 minutes, so were given the preference test again but with waxworms instead of mealworms. Of 41 individuals, 85% chose and ate a worm (either waxworm or mealworm) as their first choice, demonstrating that the birds preferred worms to seeds. For the four birds that preferred waxworms they received waxworms for all of their following experimental trials and the other birds all received mealworms. After the preference test, we gave the birds a pre-training task consisting of a 24-well tray filled with sand. We buried mealworms underneath the sand in ten randomly chosen wells, scattered ten sunflower seeds (dehusked) randomly on the surface (Fig. 1a) and recorded the first food chosen. We ran this task to confirm that the birds would forage on the tray, that the seeds were easier to access than the worms, and that the birds could not detect the buried worms either visually or through smell. 38 of the 39 birds tested chose a seed as their first choice, instead of searching in the sand, suggesting that the birds had to be trained to find the buried worms. Next, we trained the birds to forage for high-value food in sand. The purpose of this training was to teach them that when they were presented with a tray with sand, their preferred food item (i.e. worms) could be found under the sand, and in a patchy distribution. We acknowledge this training does not necessarily mean that foraging in the sand became habitual. Nevertheless, because birds became familiar with searching through the sand in the context of this novel foraging situation, we considered foraging in the sand as being their trained behaviour, and a switch to eating food on the surface of the tray was considered a plastic response. Birds were trained in a step-wise progression. In the first step, we baited all 24-wells with hidden worms, two of which were partially visible to encourage birds to search. Birds progressed to the next step if they ate five worms within one hour (n = 40). The second step was similar to the first, except only ten wells were baited (i.e. patchy distribution), one of which was partially visible. Birds progressed if they ate three worms in one hour. The final step was the same as the second but the worms were hidden in different wells compared to step two. The birds completed training if they ate three worms from this tray. Steps were repeated until birds progressed and completed the training (n = 35). Of the 41 individuals who received the food preference test, six did not complete training due to welfare concerns or time constraints.

**Figure 1.**
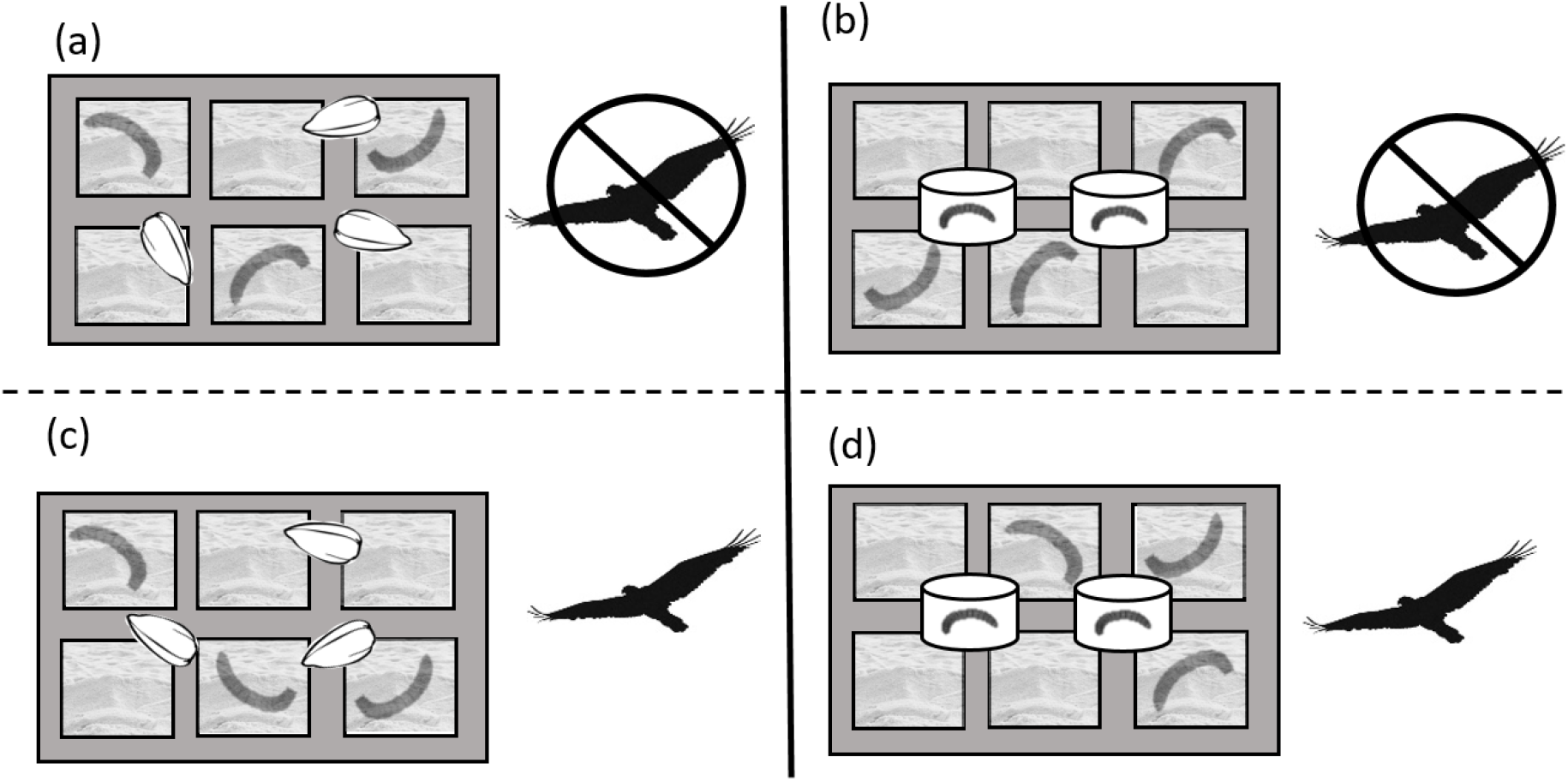
The four treatments are illustrated. In each treatment, we presented great tits with a 24-well tray filled with sand (six wells are shown here for illustrative purposes) and buried mealworms underneath the sand in ten of the 24 wells. The first treatment (a) had ten sunflower seeds (dehusked) on the surface and was presented as the pre-training task. The second (b) had two mealworms in transparent cases on the surface. The third (c) and fourth (d) treatments were as in (a) and (b) but had the addition of a simulated attack by a model sparrowhawk.

### Food choice tasks

After completion of training, all birds received four treatments in a 2 × 2 factorial design (Fig. 1). The first factor was the type of visible food and the second was the presence of a predator. In all four treatments, we placed the 24-well tray, baited with ten buried mealworms (high-value) in randomly assigned wells, on a stool in the centre of the experimental room and provided two artificial trees (1.53m H) as perches, each a metre from the stool. Visible food on the surface of the tray was one of two types: low-value (ten randomly scattered sunflower hearts), or high-value, where mealworms were encased in two transparent, sealed case (Fig. 1). They were encased for two reasons: one, to stop them burrowing in the sand and, two, to ensure that some individual variation in surface choice would likely arise when a high quality but difficult to access food item became available, otherwise all birds would have chosen the surface food. To avoid a carry-over effect of birds choosing high-value surface food leading to them by default choosing the visible option, the low-value visible food was always presented first. We assumed that the birds did not know that the worm inside the case was inaccessible, and expected the birds to attempt to get at the encased worm because it would be visible.

The first two treatments (visible low-value; visible high-value) were run in the absence of a predator and the third and fourth treatments were run in the presence of a taxidermy sparrowhawk (*Accipiter nisus*) to simulate an increased perception of predation risk (Fig. 1). Taxidermic mounts are an effective way to simulate predation risk (Carlson, Pargeter and Templeton, 2017), and have been used effectively on similar experiments in great tits (Kalb, Anger and Randler, 2019). During the third and fourth treatment, when an individual first landed on the tray to make a food choice, we released the ‘hawk’ from behind a sheet via a pulley system, to ‘fly’ across the room and ‘hide’ in a cardboard box. The order in which the visible food alternatives were presented during the two predator trials was chosen randomly to account for possible carry-over effects of the predator attack on food choice in subsequent trials.

For the four treatments, we determined all the food choices made by the birds in four minutes from video recordings. We scored food choices as ‘hidden’ (two or more pecks in the sand in the same well), or ‘visible’ (choose a seed and remove it from the tray, or touch the transparent case with foot or beak). To examine the possibility that the effects of either inhibitory control or exploration behaviour were short-lived rather than persistent, we analysed both 1) first food choice only, and 2) the proportion of visible choices out of the total number of choices made over the four minutes (henceforth, total choices). Additionally, these separate analyses were important for the visible high-value food because individuals’ choices in this experiment may have depended on their experience with the transparent casing. On their first choice, we could not assume that the birds were aware of the encased worm being inaccessible and if inhibitory control or exploration behaviour were involved in their choice, they may have required time to learn about the contingencies of this food item. Great tits sometimes flicked over the seeds with their beaks, which we did not count as a choice. A second coder (C.A.T) watched 20% of the videos to ensure the records of food choice were not biased. Strong agreement was found between raters (intraclass correlation coefficient: first choice; 100% similarity; total choices; ICC = 0.977, 95% confidence interval = 0.938-0.994).

### Statistical analysis

Data were analysed in R version 3.6.0 (R Core Team 2019). To investigate if individuals were consistent in their food choices across treatments we performed a repeatability analysis using the rptR package (Stoffel, Nakagawa and Schielzeth, 2017). Unadjusted (single variable of individual as a random effect) and adjusted (all variables contained in the model average) repeatabilities were measured for the four models mentioned below: first choice and total choices, with separate models for the effects of detour-reaching and exploration. The two unadjusted models used different datasets, due to differences in sample size. If the observed individual differences in foraging behaviour reflected intrinsic differences among individuals, then we expected the adjusted and unadjusted values to be similar. If the observed differences were caused by environmental covariation with the experimental conditions, then we expected adjusted repeatability to be lower than unadjusted values. Finally, the adjusted repeatability should be higher than the unadjusted if the experimental conditions masked among individual differences in the foraging behaviour.

For the main analyses, we used the *lme4* package (Bates *et al*., 2015) to create four models: two were based on the first choice (models 1 and 3) and two on the proportion of total choices (models 2 and 4), with either detour-reaching score (models 1 and 2, n = 29) or exploration score (models 3 and 4, n = 35) as the main explanatory variables. We included inhibitory control and exploration behaviour in separate models to avoid over-parameterisation, and because they are not correlated in this population and likely have independent effects on behaviour (Davidson G. L. unpublished data). The response variable for the first choice models was a binary ‘hidden’ (0) or ‘visible’ (1), and for the total choice models was the proportion of visible food choices out of the total number of choices made. All models were generalised linear mixed models (GLMM) with a binomial error distribution and a logit link function, with individual ID fitted as a random effect. All models had predator treatment (yes or no), visible food type (seed or encased worm), age (adult or juvenile), resident habitat (deciduous or coniferous), sex and the interaction effect between predator treatment and visible food, included as explanatory variables. We included resident habitat because habitat origin affects food choice in our populations (Serrano-Davies, O’Shea and Quinn, 2017). The results for age, habitat and sex are included in the supplementary material. Our predictions were tested by the inclusion of detour-reaching score (a proportion out of ten, treated as continuous) in models 1 and 2, and exploration score (continuous) in models 3 and 4, and their interactions with both visible food type and predation risk.

We used the *DHARMa* package (Hartig 2019) to check model fit and to test model assumptions. We used the *dredge* function from the MuMIn package (Barton 2019) and an information-theoretic approach in combination with model averaging (Grueber *et al*., 2011) to generate the models with the most support, taken from the global model. The information-theoretic approach compares multiple models (i.e. hypotheses) simultaneously and we calculated the amount of support for each model using Akaike’s Information Criterion corrected for small sample sizes (AICc) (Burnham and Anderson, 2002). Models with a ΔAICc <2 were retained as the ‘top’ models that included the most important explanatory variables. We report the averaged weighted parameter estimates across all models in the top set.

## Results

### Adjusted and unadjusted repeatabilities

Repeatability analyses confirmed that individuals differed consistently from one another in their first and total choices (Table 2). The model’s fixed effects masked the between individual differences in the first choice made because adjusted repeatability was higher when these effects were controlled (Table 2). Repeatability estimates for the total choices analyses were unaffected by whether or not the fixed effects were included.

**Table 2.** R value, p value and confidence intervals for repeatability analysis calculated in R using the rptR package. The first choice and total choices for detour reaching and exploration were analysed in two models. Unadjusted: single variable of individual as a random effect, and Adjusted: all variables from the model average. All values are from the link-scale approximation. Due to the smaller number of birds that completed the detour-reaching task than completed the exploration behaviour, the sample size for the models including detour-reaching are smaller.

| Dataset | N | Model | R value | P value | 95% Confidence interval |
| --- | --- | --- | --- | --- | --- |
| First choice |  |  |  |  |  |
| Detour reaching data subset | 29 | Unadjusted | 0.16 | 0.041 | 0; 0.37 |
|  |  | Adjusted | 0.32 | 0.01 | 0; 0.74 |
| Exploration data subset | 35 | Unadjusted | 0.22 | 0.006 | 0.02; 0.41 |
|  |  | Adjusted | 0.47 | <0.001 | 0.08; 0.84 |
| Total choices |  |  |  |  |  |
| Detour reaching data subset | 29 | Unadjusted | 0.30 | <0.001 | 0.10; 0.48 |
|  |  | Adjusted | 0.32 | <0.001 | 0.10; 0.47 |
| Exploration data subset | 35 | Unadjusted | 0.31 | <0.001 | 0.11; 0.47 |
|  |  | Adjusted | 0.27 | <0.001 | 0.10; 0.41 |

### Food choice and predation risk

Whether birds switched to the visible food depended on both the value of the visible food and the predation risk treatment. This was true for both the first choice (Tables 3 and 4; Fig. 2a) and total choices (Tables 5 and 6; Fig. 2b), and there was a similar pattern of response across both (compare Fig. 2a and 2b). Birds were more likely to switch to the visible food when it was high value, and when there was no predator present. Despite this, some birds switched to feeding on the surface even when the value of the visible food was low compared to the hidden food, and even when the predator was present, demonstrating individual variation in foraging plasticity.

**Table 3.** Analysis for first choice made by the great tits including detour-reaching score as an explanatory variable. The values shown are the average of all the top models within two AICc of the best model. A positive value for the estimate means the visible food is more likely to be selected than the hidden food. The relative importance (averaged weight: sum of Akaike weights) for each parameter is shown. ‘Age’ as a fixed effect and two interactions, ‘Detour*Visible food’ and ‘Detour*Predator’ are excluded because they did not appear in any of the top models.

| *Inhibitory control*<br>*First choice* | **Estimate** | **Stan. Error** | **95% Confidence interval** | **Averaged weight** | **P value** |
| --- | --- | --- | --- | --- | --- |
| **Intercept** | 2.31 | 0.99 | 0.37; 4.25 |  | 0.02 |
| **Fixed effects** |  |  |  |  |  |
| **Predator** |  |  |  |  |  |
| No | 0 | 0 |  |  |  |
| Yes | -4.14 | 1.03 | -6.16; -2.11 | 1.0 | <0.001 |
| **Visible food** |  |  |  |  |  |
| Encased worm | 0 | 0 |  |  |  |
| Seed | -3.01 | 0.92 | -4.80; -1.21 | 1.0 | 0.001 |
| **Detour** | -0.45 | 1.16 | -2.72; 1.81 | 0.26 | 0.70 |
| **Sex** |  |  |  |  |  |
| Female | 0 | 0 |  |  |  |
| Male | -0.18 | 0.51 | -1.19; 0.82 | 0.24 | 0.72 |
| **Habitat** |  |  |  |  |  |
| Coniferous | 0 | 0 |  |  |  |
| Deciduous | 1.67 | 0.83 | 0.05; 3.29 | 1.0 | 0.05 |
| **Interactions** |  |  |  |  |  |
| **Predator\*Visible food** | 3.68 | 1.22 | 1.28; 6.08 | 1.0 | 0.003 |

**Table 4.** Analysis for first choice made by the great tits including exploration score as an explanatory variable. The values shown are the average of all the top models within two AICc of the best model. A positive value for the estimate means the visible food is more likely to be selected than the hidden food. The relative importance (averaged weight: sum of Akaike weights) for each parameter is shown.

| *Exploration*<br>*First choice* | **Estimate** | **Stan. Error** | **95% Confidence**<br>**interval** | **Averaged**<br>**weight** | **P value** |
| --- | --- | --- | --- | --- | --- |
| **Intercept** | 0.54 | 1.07 | -1.56; 2.63 |  | 0.62 |
| **Fixed effects** |  |  |  |  |  |
| **Predator** |  |  |  |  |  |
| No | 0 | 0 |  |  |  |
| Yes | -5.21 | 1.25 | -7.66; -2.76 | 1.0 | < 0.001 |
| **Visible food** |  |  |  |  |  |
| Encased worm | 0 | 0 |  |  |  |
| Seed | -1.70 | 0.95 | -3.57; 0.17 | 1.0 | 0.08 |
| **Exploration** | 0.29 | 0.12 | 0.05; 0.53 | 1.0 | 0.02 |
| **Sex** |  |  |  |  |  |
| Female | 0 | 0 |  |  |  |
| Male | -0.12 | 0.45 | -0.99; 0.75 | 0.15 | 0.79 |
| **Age** |  |  |  |  |  |
| Adult | 0 | 0 |  |  |  |
| Juvenile | 0.23 | 0.66 | -1.06; 1.52 | 0.23 | 0.73 |
| **Habitat** |  |  |  |  |  |
| Conifer | 0 | 0 |  |  |  |
| Deciduous | 1.28 | 1.12 | -0.91; 3.48 | 0.74 | 0.26 |
| **Interactions** |  |  |  |  |  |
| **Predator\*Visible food** | 4.39 | 1.31 | 1.81; 6.96 | 1.0 | < 0.001 |
| **Exploration\*Visible food** | -0.26 | 0.12 | -0.49; -0.02 | 1.0 | 0.03 |
| **Exploration\*Predator** | 0.02 | 0.06 | -0.10; 0.14 | 0.19 | 0.72 |

**Figure 2.**
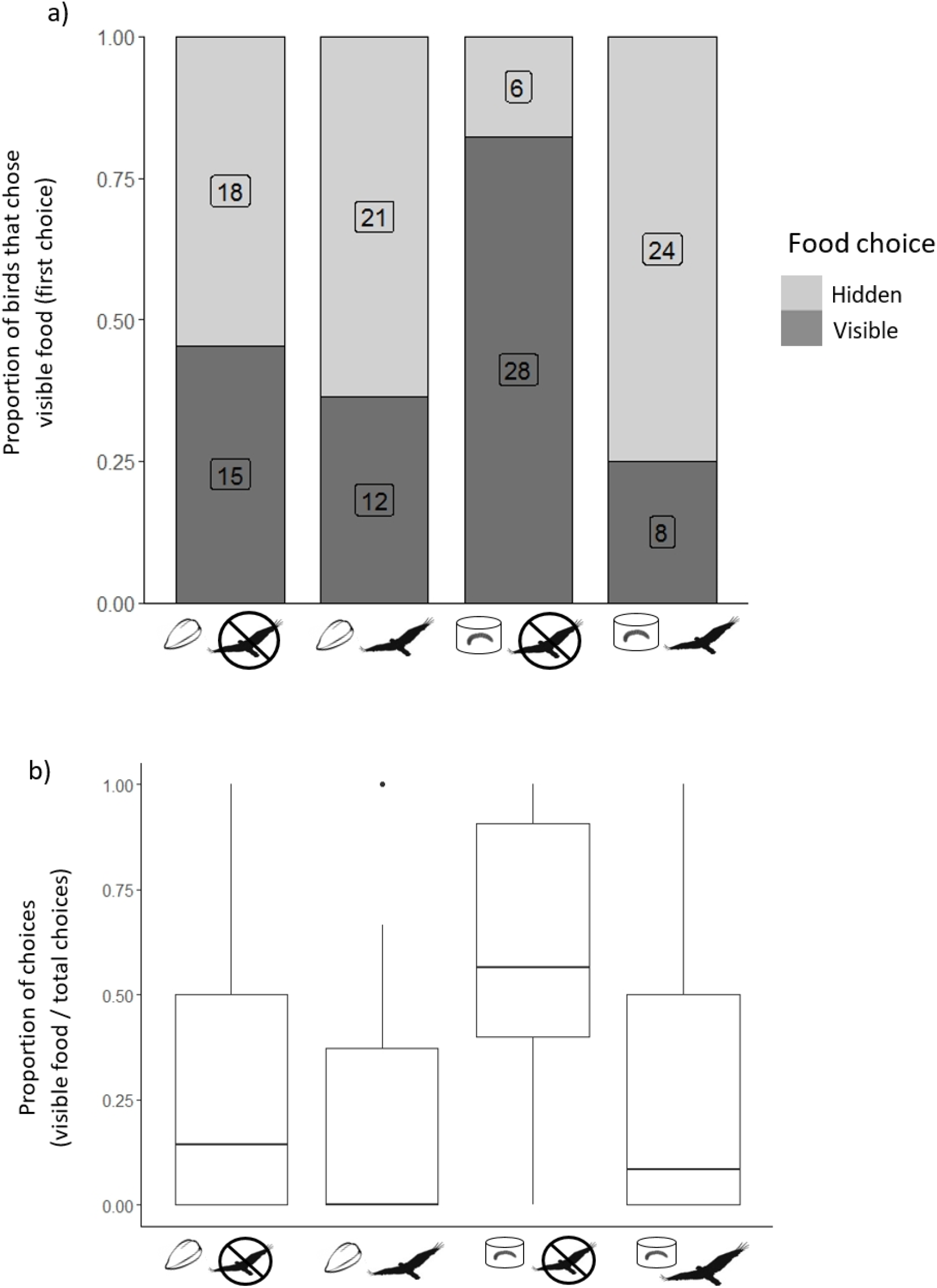
The four treatments with (a) the proportion of birds that chose the visible food and that chose the hidden food on their first choice and (b), the proportion of choices for the visible food out of the total number of choices made in four minutes. For (a), sample sizes are given on each bar and for (b) the 25^th^ and 75^th^ quartiles and median are shown and the whiskers are ±1.5*IQR.

**Table 5.** Analysis for total choices made by the great tits in four minutes including detour-reaching score as an explanatory variable. The values shown are the average of all the top models within two AICc of the best model. A positive value for the estimate means the visible food is more likely to be selected than the hidden food. ‘Age’ as a fixed effect and the interaction ‘Detour*Predator’ have been excluded because they did not appear in any of the top models. The relative importance (averaged weight: sum of Akaike weights) for each parameter is shown.

| *Inhibitory control*<br>*Total choices* | **Estimate** | **Stan. Error** | **95% Confidence interval** | **Averaged weight** | **P value** |
| --- | --- | --- | --- | --- | --- |
| **Intercept** | 0.13 | 0.97 | -1.76; 2.02 |  | 0.89 |
| **Fixed effects** |  |  |  |  |  |
| **Predator** |  |  |  |  |  |
| No | 0 | 0 |  |  |  |
| Yes | -2.41 | 0.34 | -3.07; -1.75 | 1.0 | <0.001 |
| **Visible food** |  |  |  |  |  |
| Encased worm | 0 | 0 |  |  |  |
| Seed | -0.86 | 0.48 | -1.80; 0.08 | 1.0 | 0.08 |
| **Detour** | 1.77 | 1.45 | -1.07; 4.61 | 1.0 | 0.23 |
| **Sex** |  |  |  |  |  |
| Female | 0 | 0 |  |  |  |
| Male | -0.69 | 0.75 | -2.17; 0.78 | 0.61 | 0.36 |
| **Habitat** |  |  |  |  |  |
| Coniferous | 0 | 0 |  |  |  |
| Deciduous | 0.43 | 0.62 | -0.79; 1.64 | 0.47 | 0.49 |
| **Interactions** |  |  |  |  |  |
| **Predator\*Visible food** | 2.47 | 0.50 | 1.49; 3.45 | 1.0 | <0.001 |
| **Detour\*Visible food** | -2.54 | 0.96 | -4.42; -0.66 | 1.0 | 0.009 |

**Table 6.** Analysis for total choices made by the great tits in four minutes including exploration score as an explanatory variable. The values shown are the average of all the top models within two AICc of the best model. A positive value for the estimate means the visible food is more likely to be selected than the hidden food. ‘Age’ as a fixed effect has been excluded because it did not appear in any of the top models. The relative importance (averaged weight: sum of Akaike weights) for each parameter is shown.

| *Exploration*<br>*Total choices* | **Estimate** | **Stan. Error** | **Confidence**<br>**interval** | **Averaged**<br>**weight** | **P value** |
| --- | --- | --- | --- | --- | --- |
| **Intercept** | 0.62 | 0.51 | -0.38; 1.63 |  | 0.23 |
| **Fixed effects** |  |  |  |  |  |
| **Predator** |  |  |  |  |  |
| No | 0 | 0 |  |  |  |
| Yes | -2.96 | 0.34 | -3.63; -2.29 | 1.0 | <0.001 |
| **Visible food** |  |  |  |  |  |
| Encased worm | 0 | 0 |  |  |  |
| Seed | -1.70 | 0.31 | -2.31; -1.08 | 1.0 | <0.001 |
| **Exploration** | 0.08 | 0.04 | -0.001; 0.17 | 1.0 | 0.06 |
| **Sex** |  |  |  |  |  |
| Female | 0 | 0 |  |  |  |
| Male | -1.18 | 0.45 | -2.07; -0.30 | 1.0 | 0.009 |
| **Habitat** |  |  |  |  |  |
| Conifer | 0 | 0 |  |  |  |
| Deciduous | 0.36 | 0.51 | -0.64; 1.36 | 0.48 | 0.48 |
| **Interactions** |  |  |  |  |  |
| **Predator\*Visible food** | 2.50 | 0.41 | 1.70; 3.31 | 1.0 | <0.001 |
| **Exploration\*Visible food** | -0.04 | 0.05 | -0.13; 0.05 | 0.56 | 0.41 |
| **Exploration\*Predator** | 0.10 | 0.04 | 0.03; 0.18 | 1.0 | 0.006 |

### Detour-reaching and inhibitory control

Our assay of inhibitory control, the detour reaching score, did not predict whether birds switched to the visible food in their first choice, either as a main effect, or in either of the interactions with food value or predation risk (Table 3; for weights of the top models see Table S1). An interaction between detour reaching with visible food type did predict choices in the total choice analysis (Detour*Visible food; Table 5; Fig. 3; for weights of the top models see Table S2). Birds that had a high score on the detour-reaching task were more likely to choose the visible food than birds that had a low score, and only when the visible food was high value (Tukey posthoc test: Estimate= 0.73, St. Error = 0.24, z = 3.02, *P* = 0.0025; Fig. 3). The interaction between detour-reaching score and predation risk on food choice did not appear in the average of the top models.

**Figure 3.**
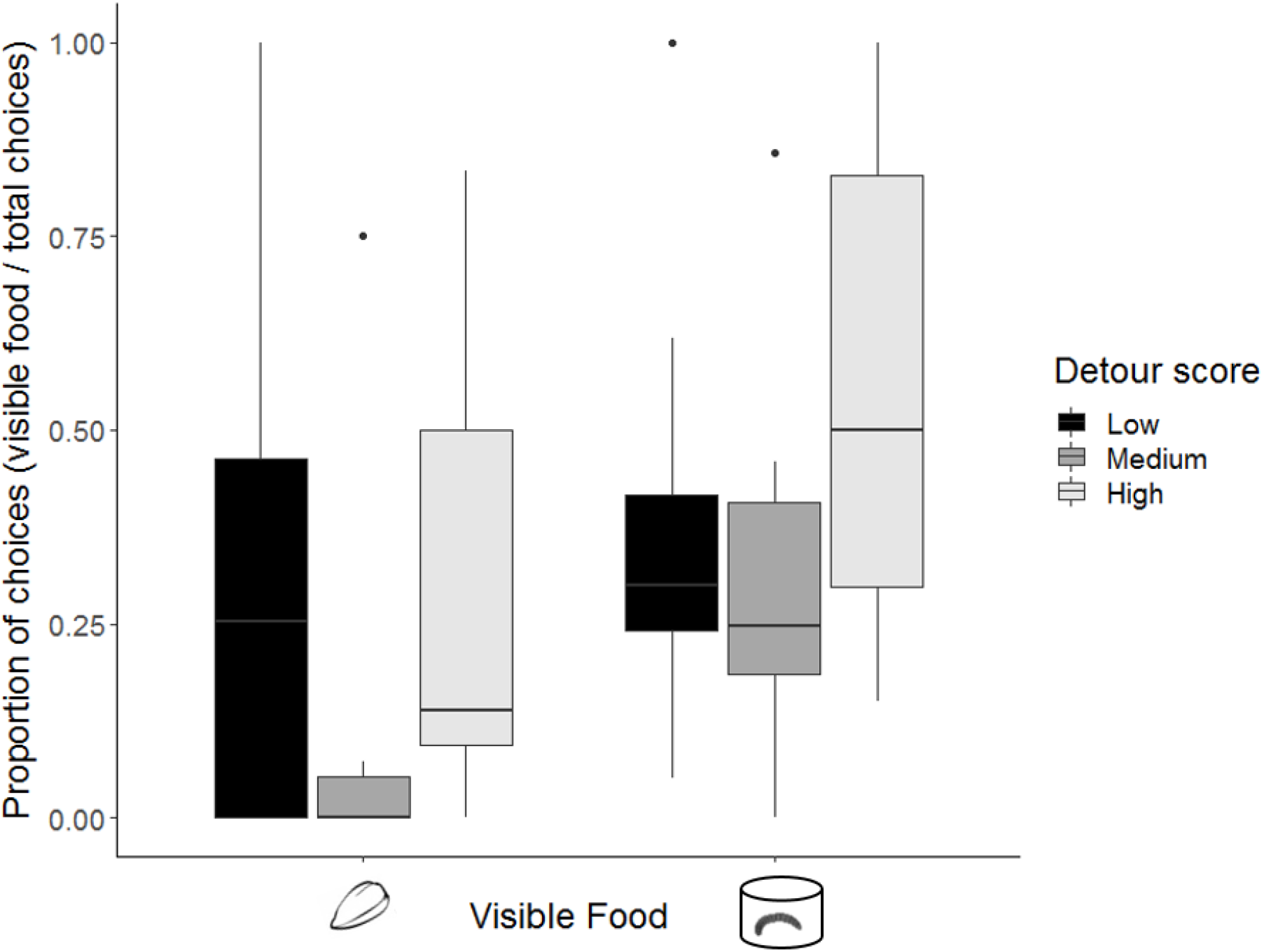
Proportion of choices for the visible food out of the total number of choices made in four minutes, against detour-reaching score, for each visible food type (averaged for the two treatments with the same visible food type). For illustrative purposes, the detour-reaching score has been split into three groups; Low, Medium and High (Range, median, mean: Low: 0 – 0.3, 0.2, 0.2, n = 10; Medium: 0.4-0.5, 0.4, n = 8, 0.41; High: 0.6-0.8, 0.7, 0.69, n = 11).

### Exploration behaviour

Exploration behaviour, a proxy of the RPPA, had a positive main effect on whether birds switched to the visible food for first choice but not total choices (Tables 4 and 6). Exploration behaviour especially influenced first choice when the visible food was high value (Exploration*Visible food; Table 4; Fig. 4; for weights of the top models see Table S3); fast explorers were more likely than slow explorers to switch to the visible food (Tukey posthoc test: Estimate: 0.12, St. Error = 0.053, z = 2.27, *P* = 0.023; this model only converged when age, sex, predator treatment and habitat were excluded). This interaction was non-significant for total choices (Exploration*Visible food; Table 6). The effect of exploration on whether birds switched from the hidden to the visible food was not affected by predation risk in the first choice analysis (Exploration*Predator, Table 4). However, in the total choice analysis, an interaction between exploration and predator treatment did influence the switch to the visible food (Exploration*Predator; Table 6; Fig. 5; for weights of the top models see Table S4). Fast explorers were more likely to choose the visible food than slow explorers, but only in the presence of a predator (Tukey posthoc test: Estimate = 0.19, St. Error = 0.06, z = 3.02, *P* = 0.003; Fig. 5).

**Figure 4.**
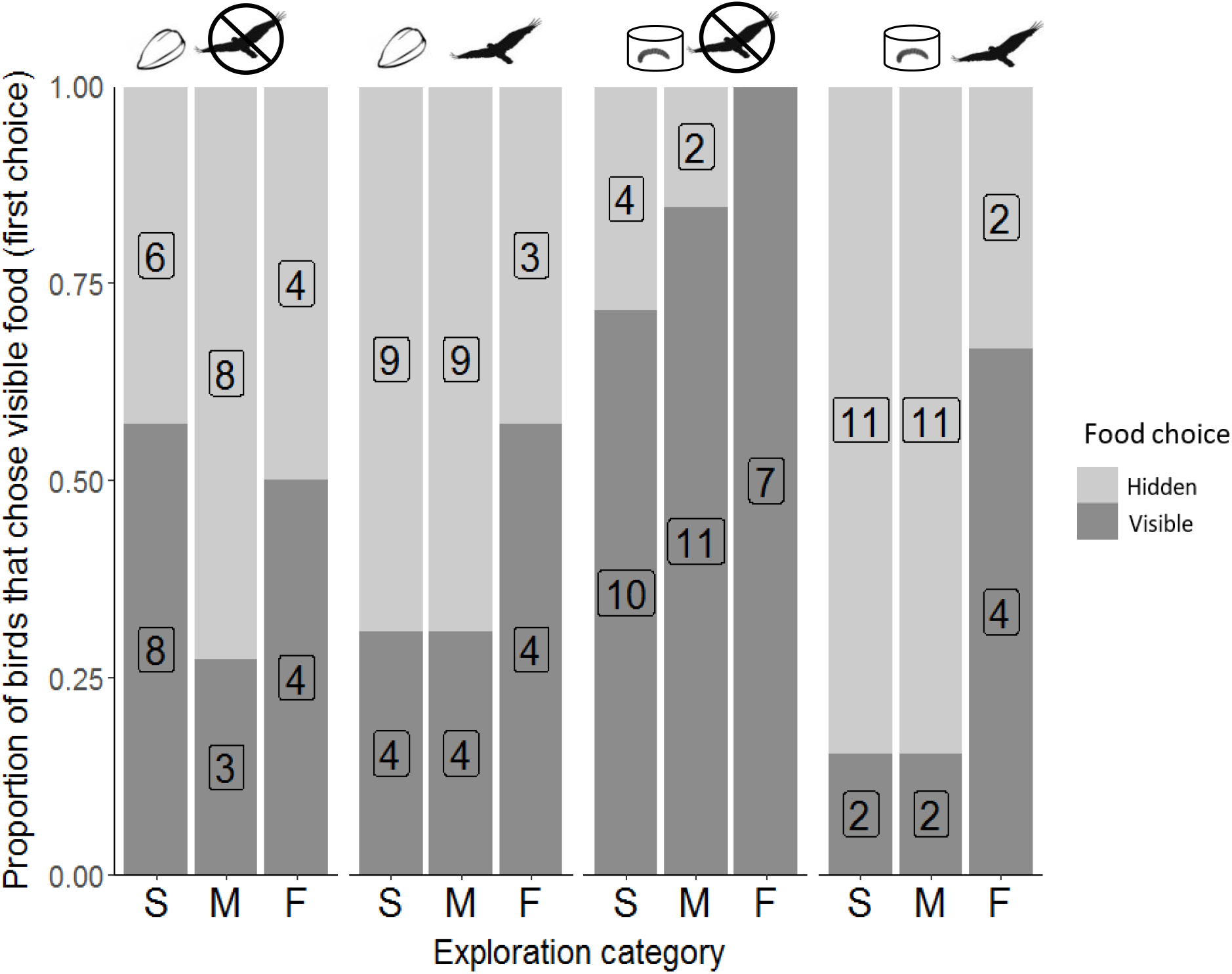
Proportion of birds that chose visible food depending on their exploration score, for first choice only. For illustrative purposes, the continuous exploration score has been split into three categories slow, medium and fast (Range, median, mean: Slow: 1-2, 1, 1.43, n = 14; Medium: 3-10, 7, 6.38, n = 13; Fast: 12-29, 15.5, 18.5, n = 8). Sample sizes are given on each bar.

**Figure 5.**
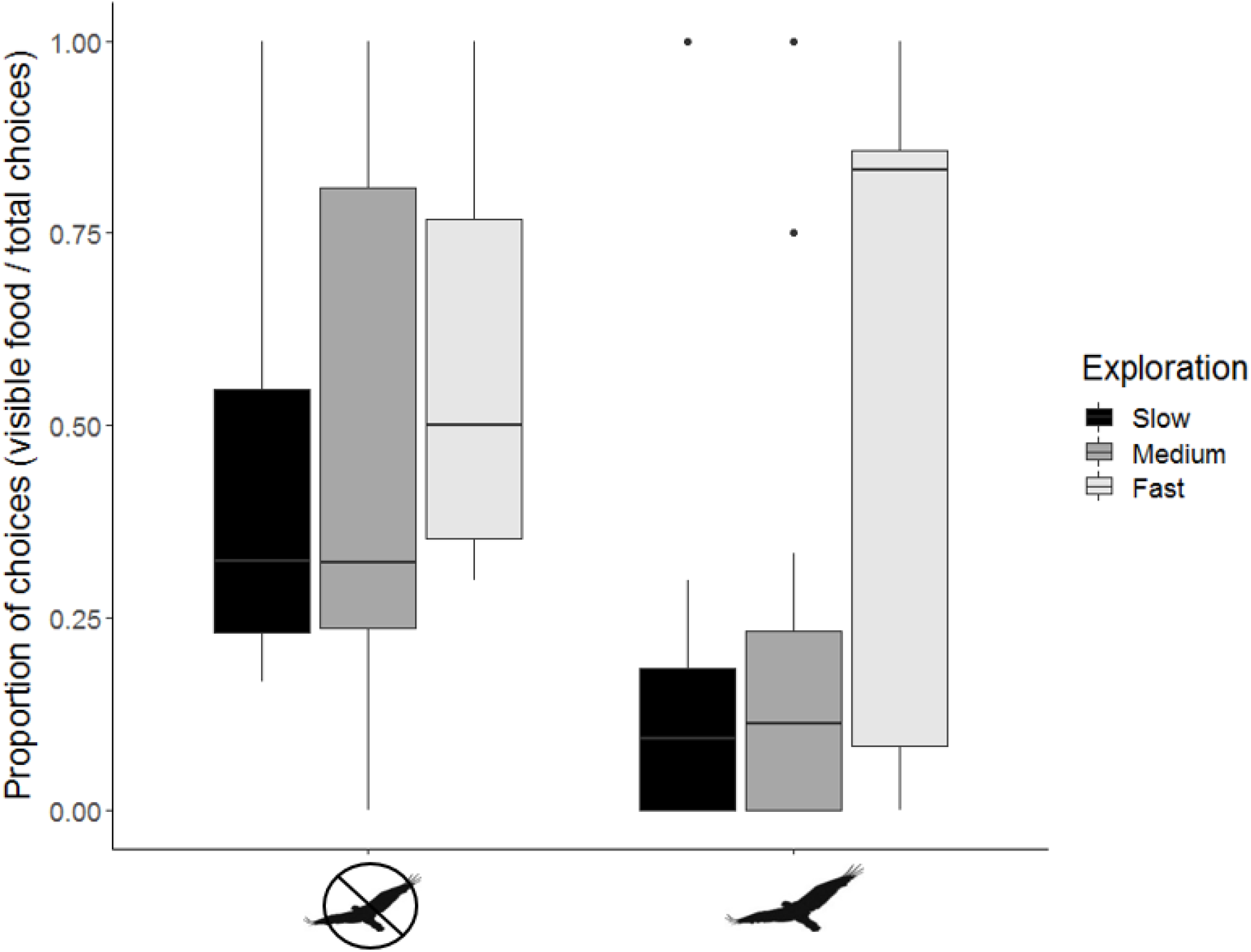
Proportion of choices for the visible food out of the total number of choices made in four minutes, against exploration score, for each predator treatment (averaged across the two food types). For illustrative purposes, the continuous exploration score has been split into three categories; slow, medium and fast (Range, median, mean: Slow: 1-2, 1, 1.43, n = 14; Medium: 3-10, 7, 6.38, n = 13; Fast: 12-29, 15.5, 18.5, n = 8).

## Discussion

There were consistent differences in individuals’ foraging behaviour. In the first choice analysis, the repeatability estimate adjusted for the two ecological factors, predation risk and surface food value, was twice as high as the unadjusted estimate. This demonstrates that failing to control for ecological variation can underestimate the potential population level consequences of this variation, although this was not true for the total choice repeatability estimate. Birds were more likely to show plasticity in their choice (to switch to the surface food) when both of the food rewards were of high value and when there was no risk of predation. Foraging plasticity was influenced by both inhibitory control and exploration behaviour, to some extent in the first choice, but especially in the total choices analysis. Fast explorers and birds with good inhibitory control were more plastic than slow explorers and birds with poor inhibitory control respectively, but only when the visible food was high-value. Fast explorers were also more plastic than slow explorers when under risk. Together these results reveal the complex interactions between foraging strategies, cognition, personality and environmental context, which we discuss in more detail below.

### Food value and predation risk

Foraging plasticity, here defined as switching from feeding on a familiar but hidden food source to an alternative visible food source, was influenced by the value of the alternative food that was available, and by predation risk. Although these were, or tended towards, significant main effects, for both the first choice and total choices, their interaction was especially important. Birds showed greater plasticity when the visible food was high value and there was no risk from a predator. These choices are consistent with optimal foraging theory, in which animals are expected to switch foraging tactics when the costs (e.g. of predation) start to outweigh the benefits of the current option (e.g. of energy gain on the patch) (MacArthur and Pianka, 1966; Milinski and Heller, 1978; Lima and Dill, 1990). In our experiment, the absence of a predator and the option of a high-value food, that seemed to be easier to access than the hidden food, meant that great tits chose to switch their food choice during this combination of treatments.

Even though great tits could not acquire the encased worm (high-value surface food) after their first attempt, they still persisted strongly for the duration of the trial. This may be because great tits are innovative and acquire food from challenging places (Aplin *et al*., 2015; Serrano-Davies, O’Shea and Quinn, 2017). As they did not know that the worm could not be accessed, and it was a desirable reward, they were willing to expend energy and time trying to acquire it. It could also be that the great tits were acquiring information about this new, unknown reward, in order to reduce their uncertainty about it, which is beneficial for survival and fitness (Stephens and Krebs, 1986; Mathot *et al*., 2012).

When there was a predator present, behavioural plasticity was supressed: most individuals foraged on the familiar, if hidden, food source. In contrast, there was no effect of risk on food choice when the low-value seeds were visible. This suggests that the great tits feel safer feeding on the familiar food, despite it taking more time to locate than the visible, surface food. Whether animals are likely to disregard high quality foods depends on risk, certainty and reward value (Holbrook and Schmitt, 1988; Mazur, 1988; Green and Myerson, 1996). In our study, the worm in the case was likely too costly to choose when there is heightened risk because it was too difficult to obtain. An alternative explanation for why individuals chose to feed on the hidden worms in the presence of a predator is that stress reduces the ability to perform goal-directed behaviour because of the inability to assess changes in food value, as seen in a study on humans (Schwabe and Wolf, 2009). As such, when great tits were in the presence of a perceived predator, perhaps they could not accurately assess the relative value of the foods and so fell back on their trained behaviour of searching in the sand for the hidden worms. Whatever the explanation, the effect of predation and food type in combination demonstrates the ecological relevance of our treatment.

### Inhibitory control

We found support for the general hypothesis that the executive cognitive function of inhibitory control influences foraging plasticity. This influence depended on the value of the visible food: individuals with a high detour-reaching score were more likely to switch from the hidden food to the visible food when both were of a similarly high value. This outcome fits the prediction that the hidden food reward was the prepotent stimulus (Table 1): individuals with good inhibitory control were able to resist the prepotent response of continuing to feed with their learned foraging technique for the hidden food, in order to feed on a visible, apparently more accessible, and therefore more immediately rewarding, food source. Birds with poor inhibitory control were less plastic in their response and therefore did not attempt to feed on the visible food, even when it was the preferred, high-value mealworm. These results suggest that individual differences in inhibitory control will influence foraging success, particularly when food differs in value and accessibility. We also predicted that predation risk could modify effects of inhibitory control on foraging plasticity because individual differences, and/or habitual behaviour, are sometimes more pronounced under stress (Suomi, 2004; Schwabe and Wolf, 2009), or because severe predation risk could over-ride any effects of inhibitory control on behaviour (Quinn and Cresswell, 2005). However, we found no interaction between predation risk and inhibitory control, suggesting that the functional significance of this executive cognitive function is not influenced by an immediate extrinsic stressor like predation risk, although whether this extends to other kinds of stressors remains to be determined. Taken together, our results suggest that differences in foraging niches and environmentally-determined food availability, rather than an immediate stressor like predation risk, can provide insight into individual differences in inhibitory control.

The effect of inhibitory control on foraging plasticity was observed when measuring total choices, rather than the first choice only, suggesting perhaps that the interaction with the encased worm influenced their subsequent choices. One might have expected birds with good inhibitory control to quickly realise that the visible food, though similar in value and ostensibly more obtainable, was in reality inaccessible, and to switch back to the hidden food, but we found the opposite. Because there was a trade-off between perceived accessibility (visible and on the surface) and searching time (not visible and patchy), birds with higher inhibitory control may have weighed this cost differently than birds with low inhibitory control. Another possibility is that individuals with high detour-reaching scores may also differ in their motivation for food, or be more persistent than those with low scores, either because the detour-reaching task measured these traits (eg. van Horik *et al*., 2018), or because these traits co-vary with inhibitory control. A further possible explanation for birds with high inhibitory control continuing to peck at the inaccessible encased worm may be due to carry-over effects from the detour task to the food choice tasks, for example, learning that food could be accessed from the side, despite a barrier. We note that although the validity of the detour-reaching task as a measure for inhibitory control has been questioned (van Horik *et al*., 2018), we chose to use it because it remains a widely used approach, and no assay of putative underlying cognitive processes is without its limitations. Additionally, we measured success/failure on a per-trial basis, repeated ten times (as opposed to counting the number of pecks on a barrier over four trials (van Horik *et al*., 2018)), and found our measure of inhibitory control to be robust against a similar task performed in the wild (Davidson G. L. unpublished data, preliminary analysis available on request).

### Personality

We found a positive main effect of exploration behaviour on the choice for the visible food for first choice only; fast explorers were more likely than slow explorers to choose the visible food. This relationship was especially pronounced when the visible food was high value. The influence of predation risk and exploration on a choice for the visible food was not dependent on the value of the visible food. When considering choices made over the entire trial (as opposed to the first choice), fast explorers were more likely to choose the visible food, regardless of its value, under predation risk. Thus the influence of exploration behaviour on plasticity was time and predation risk dependent, but not food value dependent.

Empirical studies predict that the reactive-proactive personality axis correlates with plasticity, with some suggesting a positive relationship between plasticity and proactive personalities (information gaterhing hypothesis; Frost *et al*., 2007; Mathot *et al*., 2012; Rojas-Ferrer, Thompson and Morand-Ferron, 2019), while others suggest a negative relationship between the two (behavioural flexibility hypothesis; Verbeek, Drent and Wiepkema, 1994; Wolf, Van Doorn and Weissing, 2008; Coppens, De Boer and Koolhaas, 2010). We found that fast (proactive) explorers are more plastic, supporting the information-gathering strategy (IGS) hypothesis. Our observation that overall, birds tended to forage on the familiar food option (i.e. in the sand) when under predation risk suggests they perceived the hidden food to be a safer option, despite it being more time-consuming to acquire (even if not to consume). At least in the total choice analysis, slow individuals were unlikely to feed on the visible food source under risk of predation (Fig. 5), whereas fast individuals were more likely to prioritise the visible high value food. This also supports the pace of life syndrome hypothesis (Réale *et al*., 2010; Hall *et al*., 2015), where fast individuals prioritise immediate foraging at the risk of increased predation, and slow individuals do the opposite (Stamps, 2007; Biro and Stamps, 2008; Mazza *et al*., 2019). Moreover, if stress causes individuals to perform habitual actions (Schwabe and Wolf, 2009), perhaps slow explorers, as well as being more risk-averse (Koolhaas *et al*., 1999; Groothuis and Carere, 2005; Reale *et al*., 2007), are also affected more strongly and negatively by stress than fast individuals (Baugh *et al*., 2013), and this could be another reason that they chose the familiar option when under risk.

Our results clearly suggest that this major constraint on behavioural variation, the reactive-proactive personality axis, had an effect on foraging plasticity in our experimental setup. Several studies have found personality to have different effects in different contexts (Frost *et al*., 2007; Sih and Del Giudice, 2012). In our experiment, the association between our measure of personality on behavioural plasticity was context-dependent, but the timescale in which the behaviour was expressed was also an important variable for detecting these context-dependent responses. The value of the visible food was important in the first choice, and the presence of a predator was important for total choices, which we speculate could be related to the first choice representing sampling behaviour and their total choices over four minutes representing their average choice.

Although we previously demonstrated that exploration behaviour is repeatable in our study population (O’Shea, Serrano-Davies and Quinn, 2017), and many have shown it is also heritable (e.g. (Quinn *et al*., 2009), simultaneous repeat measures of exploration score and of foraging success (or indeed of inhibitory control and foraging success), would be necessary to establish whether correlations between these pairs of behaviour occur at the between-individual level, and really do constrain plasticity (Dingemanse and Dochtermann, 2013). Estimating behavioural covariance is challenging in general, and two factors make this especially impractical in the context of this study. One is that the sample sizes would be prohibitory, not just because they are particularly high when measuring covariation (Dingemanse and Dochtermann, 2013), but also because here the covariation occurred in the context of an interaction. Another is that arguably it would be unethical to do so, since the repeat measures would have to be separated by lengthy periods of time for them to reflect anything other than temporary environmental effects. Despite this limitation in our approach, our results demonstrate that constraints on plasticity caused by behavioural mechanisms like the RPPA are likely important, if difficult to detect.

## Conclusion

Individual variation in behavioural plasticity is an important mechanism facilitating adaptation to ecological or environmental change. Our results show substantial variation in foraging plasticity, and suggest that individual differences in cognition and personality both play context-dependent roles, that are nevertheless independent of one another. We emphasise that the population level consequences of behavioural variation may only be revealed in the light of very specific ecological conditions or gradients experienced by individuals, but that very large sample sizes are going to be needed to demonstrate phenotypic or genotypic covariance among behavioural traits.

## Supporting information

Coomes Inhibitory control, personality and foraging plasticity Supplementary Material

## Author contributions

JRC, IGK, GLD and JQ designed the study, JRC and GLD collected the data with help from IGK, JRC analysed the data with help from MSR and CAT. JRC, JQ, MSR and GLD wrote the manuscript with input from IGK. All authors approve the final manuscript and agree to be held accountable for the contents.

## Acknowledgements

We would like to thank Ivan de la Hera Fernandez and Sam Bayley for catching the great tits and transporting them to the aviary and Karen Cogan for also helping with transport. We would like to thank Luke Harman and Alan Whitaker for help setting up the experiment in the aviary. We would like to thank the landowners for permission to catch the great tits at our field sites.

## Ethics statement

This study was conducted under licences from the Health Products Regulatory Authority (AE19130_P017), the National Parks and Wildlife Service (C02/2018) and the British Trust for Ornithology. The research project received ethical approval from the Animal Welfare Body at University College Cork, and was in accordance with the ASAB (Association for the Study of Animal Behaviour) Guidelines for the Treatment of Animals in Behavioural Research and Teaching.

## Data accessibility

R Code and Data are available as supplementary material, and will be uploaded to Dryad upon acceptance.

