## Supplementary material for "Inhibitory control, personality, and manipulated ecological conditions influence foraging plasticity in the great tit": Coomes Inhibitory control, personality and foraging plasticity Supplementary Material

**Supplementary information: Inhibitory control, personality and changing ecological conditions influencing foraging plasticity in the great tit**

Results

Summary of choices

Great tits chose to search in the sand more when there was a predator present (64.3% of first choices were searching) as opposed to when there was not (34.3% of first choices were searching). When the seeds were the visible food on the surface, great tits chose to search in the sand more (55.7%) than when the case was on the surface (42.9%).

The average number of choices made when there was no predator present was lower (6.90) than when there was a predator present (5.79). When the seeds were on the surface, less choices were made on average (5.69) than when the case was on the surface (7.03). The minimum number of choices made across all trials was 1 and the maximum number of choices made across all trials was 24.

For an example of a great tit choosing each surface food type, see the video clips. https://figshare.com/s/dd27c6b7b48990e635fc

Sex, age and habitat

Our analyses did not find an effect of age, or a clear effect of sex or habitat on food choice. There was a slight tendency for females to choose the visible food more than males, however we only found this effect in one of the four models (Table 4). There was a slight tendency for birds from deciduous habitats to choose the visible food more than birds from the coniferous habitats, but we only found this effect in one of the four models (Table 1). Our analysis found no effect of age on the food chosen.

**Tables**

Table S1. First choice analysis (binary response) including detour-reaching score as an explanatory variable. The table shows the included variables, degrees of freedom (Df), AICc, delta AIC and Akaike weights (ω_i_) for the top set of models within 2 AICc of the best model.

| First choice including detour-reaching | Df | AICc | Δ AIC | ω_i_ |
| --- | --- | --- | --- | --- |
| Predator*Visible Food + Visible Food + Predator + Habitat | 6 | 124.21 | 0.00 | 0.49 |
| Predator*Visible Food + Detour + Visible Food + Predator + Habitat | 7 | 125.45 | 1.24 | 0.26 |
| Predator*Visible Food + Visible Food + Predator + Habitat + Sex | 7 | 125.60 | 1.39 | 0.24 |

Table S2. Total choice analysis (response variable as the proportion of choices of the visible food out of the total number of choices) including detour-reaching score as an explanatory variable. The table shows the included variables, degrees of freedom (Df), AICc, delta AIC and Akaike weights (ω_i_) for the top set of models within 2 AICc of the best model.

| Total choice including detour-reaching | Df | AICc | Δ AIC | ω_i_ |
| --- | --- | --- | --- | --- |
| Predator*Visible Food + Detour*Visible Food + Detour + Visible Food + Predator + Sex | 8 | 323.79 | 0.00 | 0.33 |
| Predator*Visible Food + Detour*Visible Food + Detour + Visible Food + Predator + Habitat + Sex | 9 | 324.12 | 0.33 | 0.28 |
| Predator*Visible Food + Detour*Visible Food + Detour + Visible Food + Predator | 7 | 324.73 | 0.94 | 0.20 |
| Predator*Visible Food + Detour*Visible Food + Detour + Visible Food + Predator + Habitat | 8 | 324.84 | 1.04 | 0.19 |

Table S3. First choice analysis (binary response) including exploration score as an explanatory variable. The table shows the included variables, degrees of freedom (Df), AICc, delta AIC and Akaike weights (ω_i_) for the top set of models within 2 AICc of the best model.

| First choice including exploration | Df | AICc | Δ AIC | ω_i_ |
| --- | --- | --- | --- | --- |
| Predator*Visible Food + Exploration*Visible Food + Visible Food + Predator + Exploration + Habitat | 8 | 142.67 | 0.00 | 0.29 |
| Predator*Visible Food + Exploration*Visible Food  + Exploration*Predator + Visible Food + Predator + Exploration  + Habitat | 9 | 143.51 | 0.84 | 0.19 |
| Predator*Visible Food + Exploration*Visible Food + Visible Food + Predator + Exploration + Habitat + Sex | 9 | 144.08 | 1.41 | 0.15 |
| Predator*Visible Food + Exploration*Visible Food + Visible Food + Predator + Exploration | 7 | 144.19 | 1.52 | 0.14 |
| Predator*Visible Food + Exploration*Visible Food + Visible Food + Predator + Exploration + Age | 8 | 144.50 | 1.83 | 0.12 |
| Predator*Visible Food + Exploration*Visible Food + Visible Food + Predator + Exploration + Habitat + Age | 9 | 144.60 | 1.93 | 0.11 |

Table S4. Total choice analysis (response variable as the proportion of choices of the visible food out of the total number of choices) including exploration score as an explanatory variable. The table shows the included variables, degrees of freedom (Df), AICc, delta AIC and Akaike weights (ω_i_) for the top set of models within 2 AICc of the best model.

| Total choice including exploration | Df | AICc | Δ AIC | ω_i_ |
| --- | --- | --- | --- | --- |
| Predator*Visible Food + Exploration*Visible Food + Exploration*Predator + Exploration + Visible Food + Predator + Sex | 9 | 386.66 | 0.00 | 0.29 |
| Predator*Visible Food + Exploration*Visible Food + Exploration*Predator + Exploration + Visible Food + Predator + Habitat + Sex | 10 | 386.81 | 0.16 | 0.27 |
| Predator*Visible Food + Exploration*Predator + Exploration + Visible Food + Predator + Sex | 8 | 387.16 | 0.50 | 0.23 |
| Predator*Visible Food + Exploration*Predator + Exploration + Visible Food + Predator + Habitat + Sex | 9 | 387.34 | 0.69 | 0.21 |
